# The interaction of curiosity and reward on long-term memory in younger and older adults

**DOI:** 10.1101/2020.12.14.422735

**Authors:** Liyana T. Swirsky, Audrey Shulman, Julia Spaniol

**Author notes:** Correspondence concerning this article should be addressed to Liyana T. Swirsky, Department of Psychology, 350 Victoria Street, Ryerson University, Toronto, ON M5B 1K3.

## Abstract

Long-term memory is sensitive to both intrinsic and extrinsic motivation, but little is known about the relative influence of these two sources of motivation on memory performance across the adult lifespan. The study examined the effects of extrinsic motivation, manipulated via monetary reward, and curiosity, a form of intrinsic motivation, on long-term memory in healthy younger and older adults. During the incidental encoding phase on Day 1, 60 younger and 53 older participants viewed high- and low-curiosity trivia items as well as unrelated face stimuli. Half of the participants in each age group received financial rewards for correctly guessing trivia answers. On Day 2, participants completed a trivia recall test and an old-new recognition test for the face stimuli. Both curiosity and reward were associated with enhanced trivia recall, but the effects were interactive, such that only low-curiosity items benefitted from monetary reward. Neither curiosity nor reward affected face recognition performance in either age group. These findings indicate that individual and joint effects of intrinsic and extrinsic motivation on long-term memory are relatively preserved in healthy aging, a finding that highlights the viability of motivational strategies for memory enhancement into old age. Identifying conditions under which memory for unrelated information benefits from motivational spillover effects in younger and older adults is a priority for future research.

## The interaction of curiosity and reward on long-term memory in younger and older adults

Why are some facts easily retained while others are forgotten? One contributing factor is epistemic curiosity, the intrinsic motivation to acquire information for its own sake rather than its instrumental utility (Berlyne, 1965; Loewenstein, 1994). Acute states of epistemic curiosity can influence encoding and retrieval. Much as we are likely to remember information associated with extrinsic reward (e.g., money; Adcock et al., 2006), we are also more likely to remember information that makes us curious (Kang et al., 2009). Beyond enhancing memory for interesting tidbits, curiosity may also have broad effects that extend to memory for unrelated information encountered in close temporal proximity to interesting facts (Gruber et al., 2014).

While research on the link between curiosity and memory in older adults is still scarce, existing studies indicate that older adults remain sensitive to curiosity-based memory enhancement (Galli et al., 2018; McGillivray, 2015). An open question, however, is how curiosity interacts with extrinsic incentives for learning and memory in older adults. Studies with younger adults have shown that extrinsic reward boosts memory for information that arouses little curiosity, but not for information that people are naturally curious about (Murayama et al., 2010; Murayama & Kuhbandner, 2011). Knowledge about interactions between curiosity and reward may be relevant for optimizing older adults’ cognitive performance in domains that engage both intrinsic and extrinsic motivation (e.g., continuing education; Kim & Merriam, 2010).

## Curiosity and memory in younger and older adults

Several studies have used trivia paradigms to investigate the effect of curiosity on memory for interesting information (Duan et al., 2020; Kang et al., 2009; Marvin & Shohamy, 2016; McGillivray et al., 2015; Murayama & Kuhbandner, 2011) and temporally-contiguous, unrelated information (Galli et al., 2018; Gruber et al., 2014; Stare et al., 2018). In trivia paradigms, participants first complete an incidental encoding task (but see Duan et al., 2020, for an intentional version) in which they encounter high- and low-curiosity trivia items. High and low-curiosity designations are established either through a pre-screening for each participant (e.g., Gruber et al., 2014) or based on normed ratings of trivia databases (e.g., Murayama & Kuhbandner, 2011). In some studies (Galli et al., 2018; Gruber et al., 2014; Murphy et al., 2020; Stare et al., 2018), irrelevant stimuli, such as pictures of faces, are presented during the interval between the presentation of the trivia question and the answer, when curiosity is assumed to be at peak. Then, after some delay, the researcher tests participants’ recall of high- and low-curiosity trivia answers. In studies with irrelevant information included, participants also complete a recognition task for the task-unrelated stimuli.

Studies using trivia paradigms have reported significantly higher recall of high-curiosity answers compared to low-curiosity answers in both younger adults (Duan et al., 2020; Kang et al., 2009; Marvin & Shohamy, 2016; Murayama & Kuhbandner, 2011) and older adults (Galli et al., 2018; McGillivray et al., 2015). A curiosity-driven benefit has also been observed for recognition of incidental information (e.g., face stimuli) in younger adults (Gruber et al., 2014; Stare et al., 2018) and in youth (Fandakova & Gruber, 2020), such that faces associated with high-curiosity trivia items are better recognized than those associated with low-curiosity items. One study also demonstrated this effect in older adults (Galli et al., 2018; Exp 1), but failed to replicate this finding in a second experiment (Galli et al., 2018; Exp 2). In summary, there is a reliable effect of curiosity on memory of interesting information, but the effect on temporally-contiguous irrelevant information is less well established, particularly in older adults.

## Mechanisms of curiosity-based memory enhancement

Theoretically, intrinsic incentives (e.g., curiosity, satisfaction) are derived internally, while extrinsic incentives (e.g., money, social rewards) are found in our environment. Despite this theoretical distinction, the beneficial effect of curiosity on memory has been likened to that of extrinsic reward. For younger and older adults, extrinsic motivation is linked to better memory for items associated with high point value (Castel et al., 2011; Castel et al., 2016), items encoded during anticipation of financial reward (Spaniol et al., 2014), and items followed by monetary reward feedback (Mather & Schoeke, 2011). The putative mechanism for reward-enhanced memory is dopaminergic modulation of the hippocampus (Adcock et al., 2006; Bowen et al., 2020; Wittmann et al., 2005). Studies investigating the effect of curiosity on memory have implicated many of the same neural substrates as studies investigating the effect of reward on memory, including hippocampus, dopaminergic midbrain and striatal regions (e.g., substantia nigra/ventral tegmental area and nucleus accumbens; Kang et al., 2009; Gruber et al., 2014). This overlap suggests a common neural mechanism underlying intrinsic and extrinsic motivation for supporting memory processes. However, the neural mechanisms of curiosity-motivated memory are still under debate (Gruber et al., 2019). Despite similarities in neural activity underlying curiosity and reward-motivated memory when tested separately, recent evidence suggests that intrinsic curiosity and extrinsic reward rely on distinct neural systems and have unique temporal dynamics (Duan et al., 2020).

In addition to divergence in the neural basis of memory enhancement from curiosity and reward, their cognitive mechanisms may also be distinct. For example, a recent perspective suggests that curiosity is better understood as a metacognitive state than as a reward-based mechanism (Metcalfe et al., 2020). Additional evidence for a divergence of curiosity and reward effects on cognition comes from studies of motivated attention. Whereas curiosity appears to broaden attention, leading to enhanced memory for temporally contiguous information (Gruber et al., 2014), reward tends to narrow attention, increasing the selectivity with which target information is remembered (Braver et al., 2014; Chiew & Braver, 2011). When presented alongside target information, task-irrelevant information is better suppressed when extrinsic motivational incentives are available (e.g., monetary gain, Williams et al., 2017; high point value, Hennessee et al., 2018). This reward-related selectivity effect has been demonstrated in both younger and older adults (for a review, see Swirsky & Spaniol, 2019).

Taken together, both curiosity and reward are associated with enhanced memory for target information, but reward appears to narrow attention and subsequent memory (Mather & Sutherland, 2011), whereas curiosity appears to benefit encoding of any information encountered alongside the information of interest (e.g., irrelevant faces encoded between interesting trivia in Gruber et al., 2014). Evidence for distinct neural recruitment from curiosity and reward (Duan et al., 2020) along with differences in putative cognitive mechanisms (i.e., narrowing versus broadening of memory encoding) create part of the rationale for investigating how intrinsic and extrinsic motivation influence memory in younger and older adults in the current study.

## Interactions of curiosity and reward on memory

A final consideration relevant to the current study is the interaction between intrinsic curiosity and extrinsic reward. According to self-determination theory, motivation from extrinsic sources can undermine the benefits of intrinsic motivation on learning (Deci et al., 1999; Ryan & Deci, 2000). For example, younger adults show reduced willingness to voluntarily engage in a task when performance-contingent incentives are offered for performing the task (Murayama et al., 2010). A phenomenon similar to the undermining effect has also been demonstrated in the context of a curiosity-inducing trivia paradigm. Murayam and Kuhbandner (2011) demonstrated an interaction between intrinsic curiosity and extrinsic reward in which benefits from financial reward were less effective when participants were inherently interested in the information. In this study with younger adults, half of the participants completed a typical trivia paradigm as described above, while the other half completed the same task but with monetary reward at stake for correctly guessing trivia items during encoding. Results showed that financial reward and curiosity had interactive effects, such that reward enhanced memory for low-curiosity items, but not for high-curiosity items (Murayama & Kuhbandner, 2011). Contrary to these findings, one recent study (Duan et al, 2020) failed to replicate this interaction using a similar paradigm with younger adults. Here, memory for trivia showed additive, rather than interactive, effects of financial reward and curiosity. However, there were critical differences between the paradigm used by Duan et al. (2020) and Murayama and Kuhbandner (2011) that could influence the emergence of the interaction, including the type of encoding (intentional vs. incidental) and the nature (within-subjects vs. between-subjects) and time (retrieval vs. encoding) of the extrinsic reward manipulation.

## Age differences in motivation

The interactive effect of reward and curiosity has not yet been documented in older adults. However, normative changes in dopaminergic neurotransmission as well as changes in motivational orientation suggest that this interaction may manifest differently in older adults. Compared to younger adults, older adults may be less sensitive to reward-based influences on learning (Eppinger et al., 2012) which is consistent with the well-documented age-related decline in dopaminergic and serotonergic transmission (Bäckman et al., 2010; Eppinger et al., 2011).

This reduced sensitivity to financial incentive is also consistent with theories of changes in motivation across the lifespan. According to the socioemotional selectivity theory (Carstensen, 1992; Carstensen et al., 1999), shortened time horizons cause older adults to have a present-oriented focus on emotion regulation and well-being, whereas younger adults have a future-oriented focus on exploration and knowledge acquisition. Similarly, the selective engagement theory (Hess, 2014) suggests that older adults conserve resources for tasks that they deem meaningful. As such, arbitrary financial incentives may not be sufficient to motivate older adults who are no longer focused on accruing resources for the future and who may not find the task itself meaningful.

For example, older adults’ job satisfaction is a function of their own job contributions, while younger adults’ job satisfaction is predicted by their financial rewards (Kollmann et al., 2020). Interestingly, older adults actually report less job satisfaction when they feel they are being “overpaid” (i.e., compensation exceeds perceived contributions) whereas the opposite is true for younger adults (i.e., less satisfied when perceived contributions exceed compensation; Kollmann et al., 2020). Moreover, older adults may show less temporal discounting (Eppinger et al., 2012; but see Seaman et al., 2020) as well as reduced incentive-based modulation of attention (Williams et al., 2018). At the same time, some studies have revealed similar effects of financial reward on recognition and recall in younger and older adults (Mather & Schoeke, 2011; Spaniol et al., 2014). Despite some evidence of preserved reward sensitivity, older adults’ motivational orientation and weakened response to financial incentives suggests that intrinsic motivation may be a preferable route to memory enhancement in older adults, but also that older adults may be less likely to exhibit the interaction between these factors.

## The current study

The current study had two main objectives. We sought to replicate the curiosity-related memory benefit for interesting information and irrelevant, temporally contiguous information in younger and older adults. We also sought to investigate the interaction of intrinsic and extrinsic motivational influences on memory, which had never been examined in older adults.

Understanding the interaction can inform the design of environments in which both intrinsic and extrinsic motivators are at play. For example, older adults’ self-reported reasons for participating in continuing education include their intrinsic desire to learn, as well as their interest in social contact – an extrinsic motivator (Kim & Merriam, 2010). How do these factors shape learning outcomes in older adults? Are their influences additive or interactive? The answers to these questions could inform the design of educational programs for older learners. If extrinsic incentives such as social contact add little value when older adults are intrinsically interested in the material, then instructors could save incentives (e.g., social presence in the learning environment; Cobb, 2009) for material that elicits relatively low levels of interest and curiosity.

To address these aims, younger and older adults completed a typical trivia paradigm with irrelevant face stimuli included in the encoding task and memory tested at a ∼24-hour delay. Irrelevant faces were included to test the effect of curiosity on unrelated information. Critically, half of the participants in each age group received extrinsic rewards at encoding, whereas the other half did not. Our hypotheses were as follows. First, we expected to replicate prior findings of a positive association between trivia-related curiosity and subsequent memory for trivia answers, in both younger and older adults (Galli et al., 2018; McGillivray et al., 2015). We also expected a positive association between curiosity and subsequent memory for unrelated, temporally contiguous face stimuli (Galli et al., 2018, Exp. 1; Gruber et al., 2014). Second, based on past observations of an interactive effect of intrinsic and extrinsic motivation on memory in younger adults (Murayama & Kuhbandner, 2011; but see Duan et al., 2020), we expected a similar interaction in the current study, such that reward would boost younger adults’ memory for low-curiosity but not high-curiosity material. In line with a reduced influence of extrinsic monetary reward on cognitive performance in aging (Bäckman, et al., 2006; Schott et al., 2007), however, we predicted no interaction of curiosity and reward in older adults.

## Method

### Participants

Participants in the final sample included 60 younger adults (aged 18-35; 40 female) and 53 older adults (aged 60 or older; 33 female). The final total sample size thus approximated the a-priori sample size target (*N* = 108) determined using a power analysis with G-Power (Faul, Erdfelder, Lang, & Buchner, 2007), requiring a power of at least .95 to detect a medium-sized interactiont (*f* ≥ .25) of) of a between-subjects factor and a within-subjects factor in each age group, assuming an alpha error probability of .05 and a correlation among levels of the within-subjects factor of .50 or higher. Younger adults were recruited from the community. Older adults were recruited from the Ryerson Senior Participant Pool, a database of community-dwelling seniors. Eligibility criteria included normal or corrected-to-normal vision and absence of neurological, psychiatric, or cardiovascular conditions that might affect cognitive performance. In total, 71 younger adults were tested, but 11 were excluded from the analysis due to technical issues during Session 2 (*n* = 6), failure to return for Session 2 (*n* = 4), or insufficient low-curiosity responses during the screening task (*n* = 1). A total of 74 older adults were tested, but 21 were excluded for analysis due to technical issues during Session 2 (*n* = 2), failure to return for Session 2 (*n* = 3), insufficient low-curiosity questions from the screening task (*n* = 11), scoring below 26 on the Montreal Cognitive Assessment (MoCA; Nasreddine et al., 2005, *n* = 4), and reporting an awareness of the memory test (*n* = 1).

Participants in each age group were randomly assigned to the control condition or the reward condition. All participants received CAD 36 for participation (CAD 24 after Session 1; CAD 12 after Session 2). Participants in the reward condition received an additional performance-contingent bonus of up to CAD 12.75 after Session 1. All participants provided written informed consent, and study procedures were reviewed and approved by the Ryerson University Research Ethics Board.

### Materials

The experimental tasks were programmed in E-Prime 2.0 (Psychology Software Tools, Pittsburgh, PA) and presented on a 17-inch display. Text appeared in black 32-point Arial font against a white background. For the screening task, stimuli included 282 trivia questions drawn from various online trivia databases (see Appendix A for full list of trivia items used). For the encoding task, stimuli included a participant-specific subset of 100 trivia questions from the screening task. Based on each participants’ responses during the screening task, the 100 trivia questions used in the encoding task were classified as high-curiosity (n = 40), low-curiosity (n = 40), or known (n = 20). The encoding task also included 100 gray-scale, emotionally neutral faces (50 younger-adult and 50 older-adult faces) from the CAL/PAL Face database (Minear and Park, 2004).

For the face recognition task in Session 2, stimuli included 80 faces from the encoding task (40 each from high-curiosity and low-curiosity trials) as well as 40 new faces. Half of the faces in each of these sets were younger-adult faces and half were older-adult faces. The trivia recall task included a total of 80 trivia questions, including 40 high-curiosity questions and 40 low-curiosity questions.

### Procedure

#### Session 1

Session 1 included the screening task, paper-and-pencil questionnaires, and the encoding task. Participants in the reward condition received a bonus that was contingent on performance in the encoding task, whereas those in the control condition received no bonus.

##### Screening Task

Trivia questions were presented in random order. Each trivia question was followed by a confidence rating (“How confident are you that you know the answer?”) and a curiosity rating (“How curious are you to find out the answer?”). Both ratings used a 6-point scale. The screening task terminated once the participant had identified 40 unknown high-curiosity questions (confidence rating < 6 and curiosity rating ≥ 4), 40 unknown low-curiosity questions (confidence rating < 6 and curiosity rating ≤ 3) and 20 known questions (confidence rating = 6). See Figure 1a for a schematic of the screening task.

**Figure 1.**
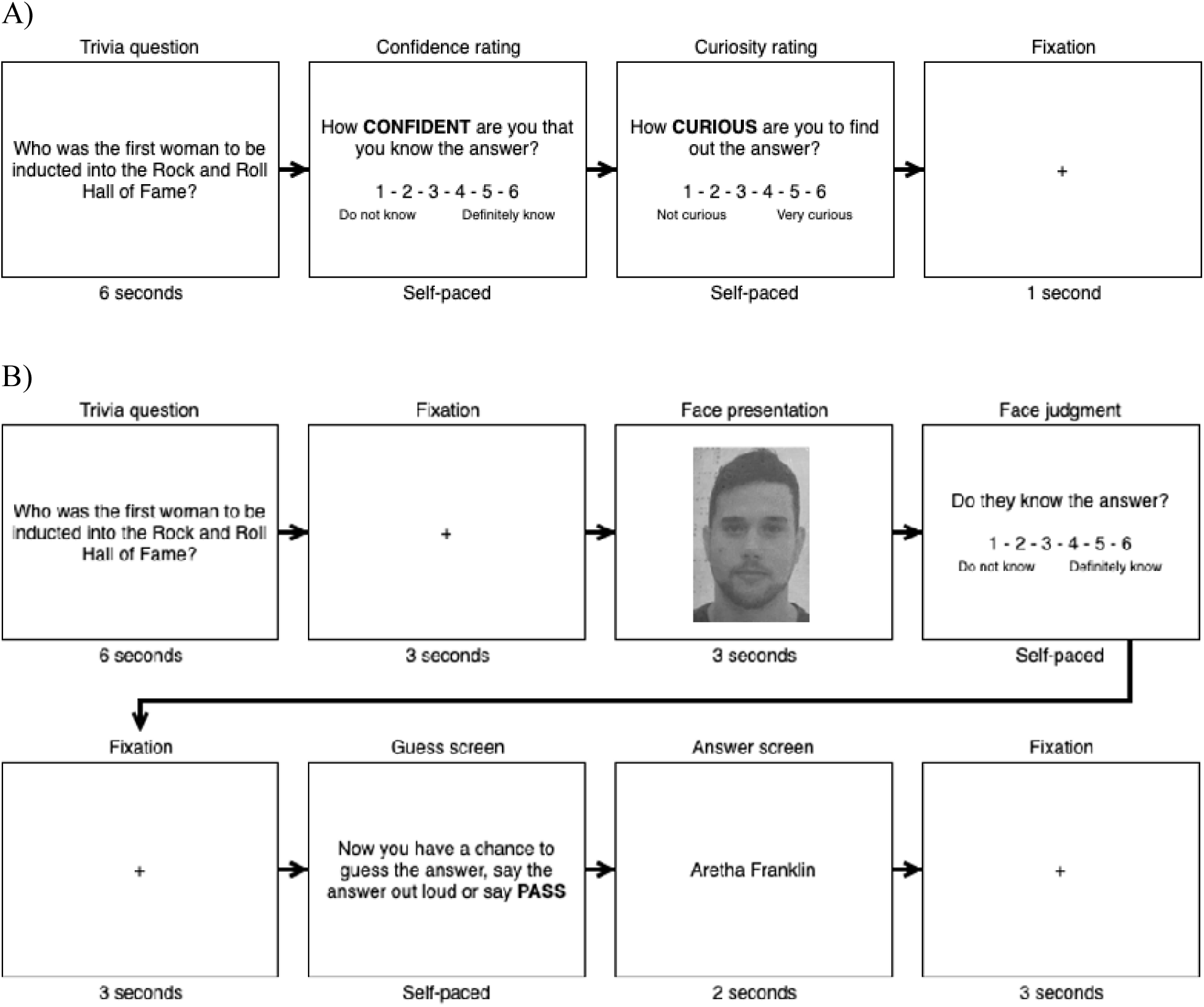
Session 1 tasks. A) Pre-screening task to determine personalized lists of high and low-curiosity trivia items for the encoding task. B) Incidental encoding task in which participants guessed answers to high and low-curiosity trivia items while making judgment about unrelated faces.

##### Encoding Task

After the participant had completed unrelated paper-and-pencil questionnaires (approximately 10 minutes), the experimenter administered the encoding task. Throughout the encoding task, participants gave verbal responses and the experimenter entered these responses using the number keys for rating screens or letter keys for trivia guesses. The responses entered by the experimenter were visible to the participant. This procedure was used to eliminate potential age differences in the demands associated with typing.

The paradigm combined elements of the encoding tasks employed by Gruber et al. (2014; incidental face stimuli) and by Murayama and Kuhbandner (2011; guessing and rewarding trivia answers). On each trial, a trivia question appeared on the screen, followed by a face stimulus and a judgment screen (“Do they know the answer?”). This judgment was included to ensure that participants attended to the task-irrelevant face. After another fixation, participants were prompted to guess the answer to the trivia question, or to pass. The final screen revealed the correct answer. Participants in the reward condition earned $0.25 per correct answer. All answers that were guessed correctly were removed from subsequent analyses. See Figure 1b for a schematic of the encoding task.

##### Background measures

After the computer tasks, participants completed the Positive and Negative Affect Schedule (PANAS; Watson et al., 1988) as well as the Need for Cognition scale (Cacioppo et al., 1984), a measure of trait-level epistemic curiosity. Older adults also completed the MoCA (Nasreddine et al., 2005) to screen for cognitive impairment.

#### Session 2

On the following day (∼24 hours post-encoding), participants returned to the laboratory and received a surprise face recognition task and trivia recall task. The order of these tasks was counterbalanced across participants within each age group and condition. The order of presentation of stimuli (trivia and faces) within both tasks was randomized for each participant. There was no difference in the tasks for those in the control condition versus the reward condition.

##### Face Recognition Task

On each trial, participants saw an old or new face stimulus and indicated, using the left and right arrow keys, whether they recognized the face from Session 1. Participants then rated their confidence in their recognition decision. See Figure 2a for a schematic of the face recognition task.

**Figure 2.**
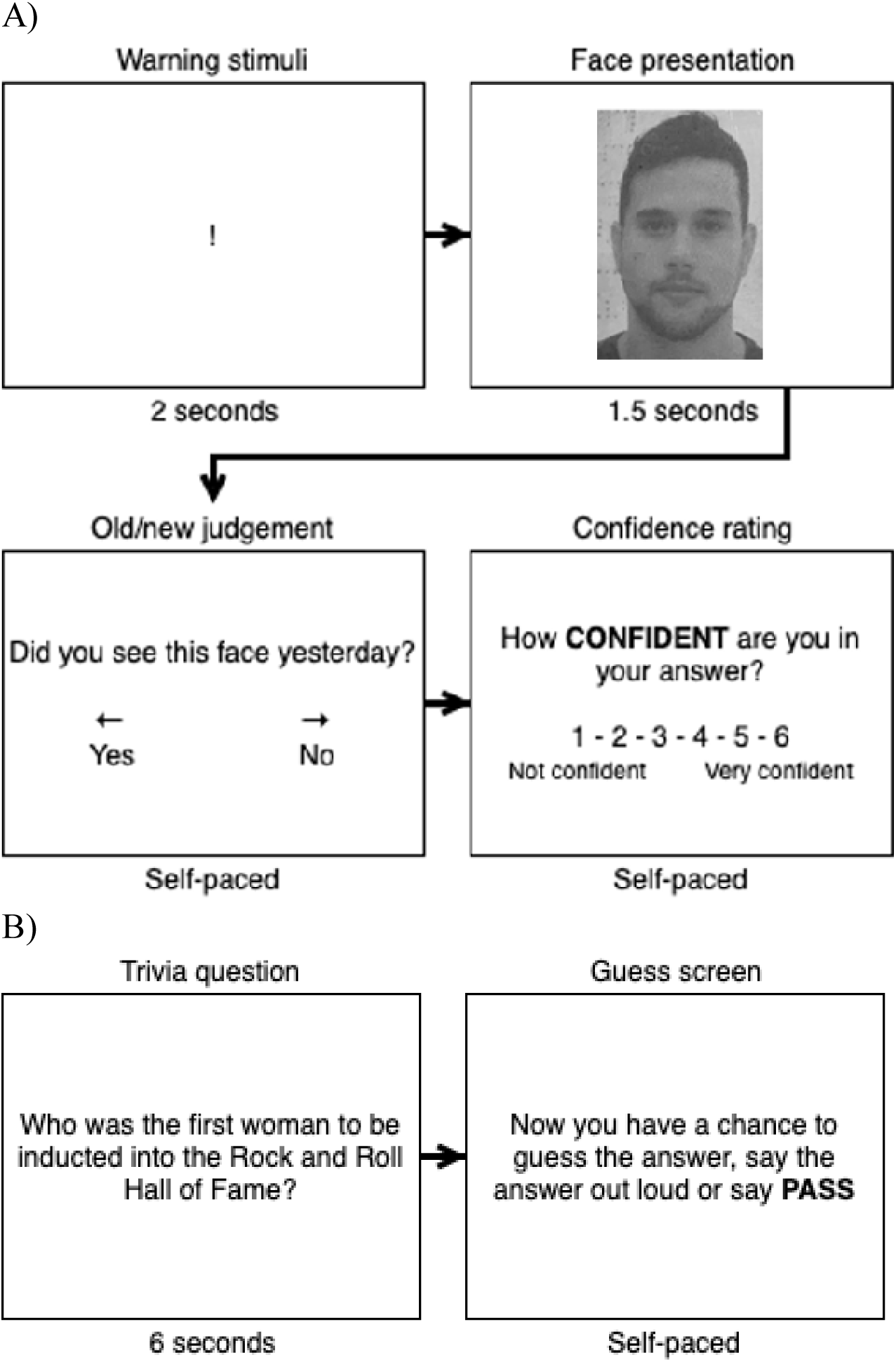
Memory tests during Session 2. The order of the tests was counterbalanced across age group and condition. A) Face recognition test in which participants identified old faces from the Session 1 encoding task. B) Trivia recall test in which participants guessed answers to trivia questions aloud while the experimenter entered their responses.

##### Trivia Recall Task

On each recall trial, participants saw a trivia question from the encoding task for 6 s, followed by a prompt to recall the answer or pass (self-paced). Participants answered verbally and the experimenter recorded their responses. See Figure 2b for a schematic of the trivia recall task.

##### Background measures

In between the two retrieval tasks, a second administration of the PANAS (Watson et al., 1988) was used to assess mood. The Shipley Vocabulary scale (Shipley, 1940) was used to assess participants’ verbal intelligence, and the Digit Symbol task (Wechsler, 1997) was used to assess processing speed.

## Results

Summary statistics for background measures are reported in Table 1. Age differences were identified with *t*-tests, also reported in Table 1. Measures with significant age differences were included as covariates in subsequent analyses. Summary statistics for curiosity ratings and trivia recall accuracy for each trivia item by age group are available in Appendix A. More detailed analyses on the relationship between trivia category and answer type with curiosity rating and recall accuracy are provided in the online supplement (see Tables S1-S4 and Figures S1-S4).

**Table 1.** Sample characteristics and age effects.

|  | Younger Adults |  |  |  | Older Adults |  |  |  | Age effects<br>*t*-value |
| --- | --- | --- | --- | --- | --- | --- | --- | --- | --- |
|  | Control |  | Reward |  | Control |  | Reward |  |  |
|  | *M* | *SD* | *M* | *SD* | *M* | *SD* | *M* | *SD* |  |
| N | 30 | — | 31 | — | 26 | — | 26 | — | — |
| N (Female) | 19 | — | 21 | — | 18 | — | 15 | — | — |
| Age, years | 25.37 | 5.49 | 24.65 | 4.84 | 69.92 | 6.87 | 70.08 | 6.20 | — |
| Age range, years | 18-35 | — | 18-35 | — | 61-86 | — | 62-89 | — | — |
| Education, years | 15.27 | 2.04 | 15.79 | 2.02 | 16.77 | 2.05 | 17.46 | 2.23 | **-4.01\*** |
| Shipley | 30.83 | 3.97 | 30.06 | 4.19 | 36.69 | 1.91 | 36.27 | 2.27 | **-10.14\*** |
| NFC | 52.47 | 5.66 | 53.58 | 5.73 | 50.00 | 10.74 | 51.35 | 6.64 | 1.80 |
| *Digit symbol* |  |  |  |  |  |  |  |  |  |
| Correct | 88.57 | 16.40 | 85.19 | 19.15 | 66.15 | 12.31 | 67.19 | 16.94 | **6.61\*** |
| Incorrect | 0.53 | 0.86 | 1.00 | 1.73 | 0.46 | 0.95 | 0.50 | 1.42 | 1.19 |
| MoCA | — | — | — | — | 28.04 | 1.31 | 28.23 | 1.07 | — |
| *Session 1 PANAS* |  |  |  |  |  |  |  |  |  |
| Positive Affect | 29.97 | 8.64 | 30.29 | 7.02 | 33.96 | 5.38 | 36.58 | 6.72 | **-3.91\*** |
| Negative Affect | 13.63 | 3.97 | 12.55 | 3.35 | 10.88 | 1.24 | 11.31 | 1.85 | **3.83\*** |
| *Session 2 PANAS* |  |  |  |  |  |  |  |  |  |
| Positive Affect | 27.93 | 9.65 | 28.74 | 9.41 | 35.54 | 7.23 | 37.27 | 7.91 | **-5.03\*** |
| Negative Affect | 12.70 | 3.53 | 12.13 | 3.02 | 11.00 | 1.44 | 10.54 | 1.45 | **3.53\*** |

*Note.* NFC = Need for Cognition; MoCA = Montreal Cognitive Assessment; PANAS = Positive Negative Affective Schedule; \* $p < .001$ . $t$ (60), two-tailed test for age group differences, equal variances not assumed.

To assess overall performance on the trivia recall task, the proportion of correctly recalled trivia answers was calculated for each participant (see Table 2 and Fig. 3b). Any trivia items that were correctly guessed during the encoding task were removed from the recall analysis. Answers were scored by two independent coders and discrepancies were resolved via discussion. Interrater reliability was near ceiling (98.3% agreement). Trial-level recall outcomes (0 = not recalled, 1 = recalled) served as the dependent variable in all subsequent mixed model analyses.

**Figure 3.**
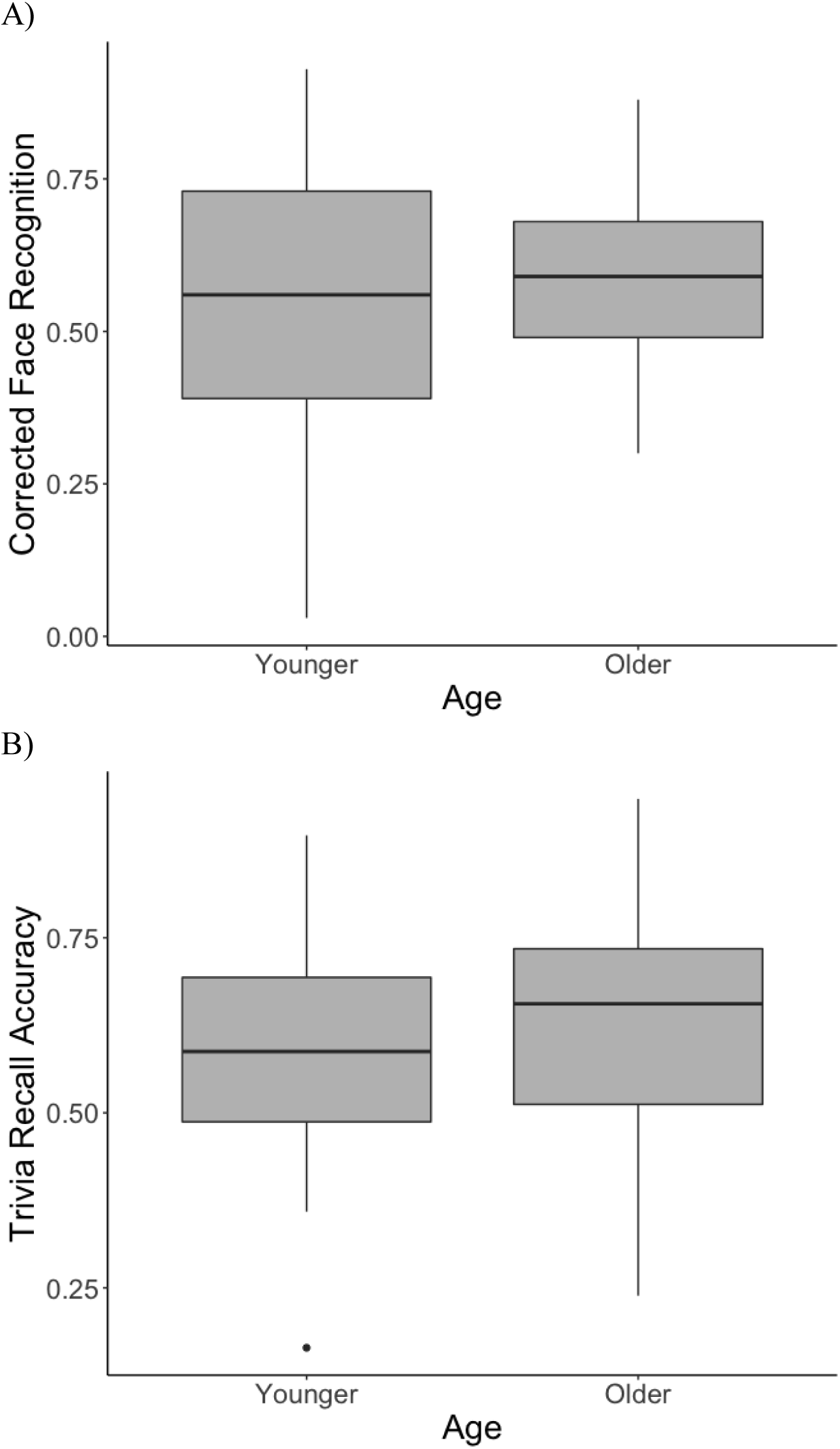
A) Older and younger adult performance on the face recognition task. B) Older and younger adult performance on the trivia recall task. Inclusion of the younger adult outlier (solid black dot) did not change the pattern of findings from the trivia recall analysis, therefore the outlier was retained in the analysis. Error bars represent the standard error of the mean.

**Table 2.**
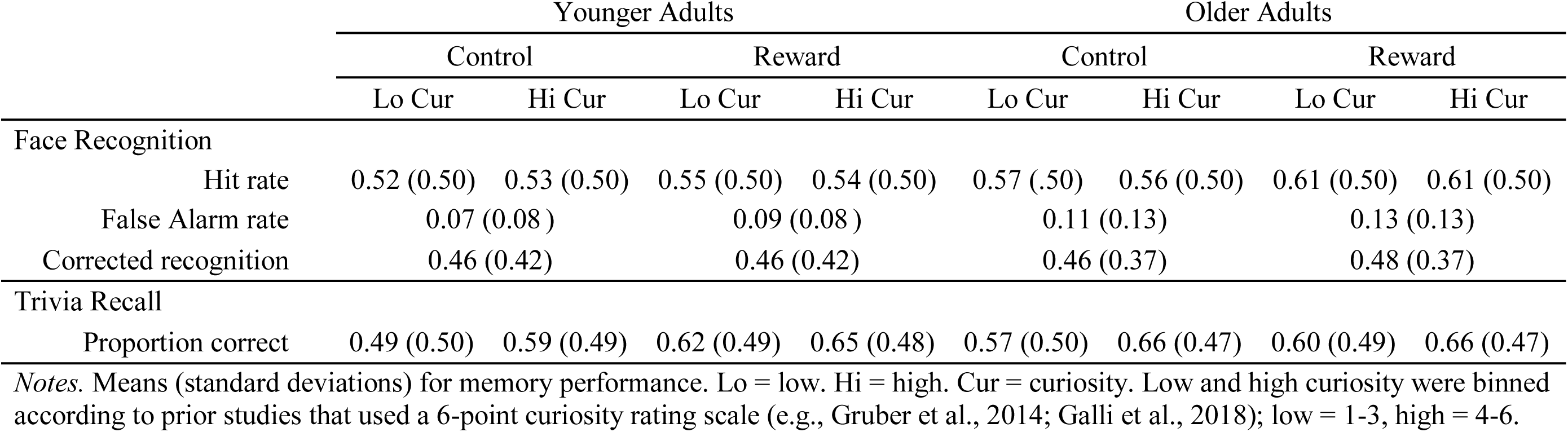
Descriptives for task performance on face recognition and trivia recall.

To assess overall face recognition accuracy, the corrected recognition rate (hit rate minus false-alarm rate) was calculated for each participant (see Table 2 and Fig. 3a). Trials in which confidence was rated as “1” were removed from the analysis to account for guessing. While accuracy provided information about overall recognition performance, only the responses to old items were informative about differences related to encoding conditions (control versus reward; high versus low curiosity). Therefore, trial recognition outcomes (0 = miss, 1 = hit) were used as the dependent variable in a series of mixed model analyses, described below.

### Mixed Model Analyses

To accommodate the nested data structure of trials within participants and the dichotomous nature of the dependent variables (trial-level recognition and recall outcomes), logistic mixed effects models were estimated for both dependent variables. All statistical analyses were carried out in R version 1.3.959 (R Core Team, 2020). Mixed models were estimated using the glmer function of the lme4 package (Bates et al., 2015), *p* values for model coefficients were estimated using the lmerTest package (Kuznetsova et al., 2017), and fixed effects and interactions were tested using the Anova function from the car package (Fox & Weisberg, 2019) and are reported as Wald chi-square tests. For ease of interpretation, fixed effects coefficients were exponentiated to calculate odds ratios (Murayama et al., 2014).

For both face recognition and trivia recall, dependent variables were regressed on curiosity rating (quasi-continuous, from 1 to 6), age group (binary), and condition (binary) as well as all possible two-way interaction terms and the three-way interaction term. Random effects models were estimated to account for repeated observations (i.e., trials) within participants. Curiosity ratings were grand mean-centered. Dichotomous variables were effect-coded to examine main effects, and dummy-coded in both directions to probe significant interactions. Levels of effect-coded predictors were assigned values of -1 or 1 to avoid the use of a reference when interpreting main effects of other variables. Because neither group was coded as zero, models with effect-coded predictors allow for the interpretation of predictor coefficients collapsing across levels of the effect-code predictor (i.e., 0 refers to an average of the -1 and 1 conditions). Levels of dummy-coded predictors were assigned values of 0 or 1 in the first instance of the variable and then the coding scheme was flipped for the second instance of the variable. With this dummy-coding approach, results can be interpreted with both levels of the variable as the reference group depending on which dummy version is included in the model, which is critical when breaking down interactions (see Table 4).

**Table 4.** Parameter estimates from the logistic mixed models used to probe the Curiosity * Condition interaction effect on trial-level trivia recall accuracy

|  | Estimate | SE | z | p | Odds ratio |
| --- | --- | --- | --- | --- | --- |
| Model 1: Condition dummy-coded with control as the reference group |  |  |  |  |  |
| Intercept | 0.36 | 0.09 | 3.81 | **<.01** | 1.43 |
| Curiosity (mean-centered) | 0.27 | 0.03 | 7.88 | **<.01** | 1.31 |
| Age group (effect-coded; ; YA: -1, OA: 1) | 0.16 | 0.09 | 1.70 | 0.09 | 1.17 |
| Condition (dummy-coded; Con: 0, Rew: 1) | 0.29 | 0.13 | 2.19 | **0.03** | 1.34 |
| Curiosity * Age Group | <0.01 | 0.03 | 0.02 | 0.98 | 1.00 |
| Curiosity * Condition | -0.11 | 0.05 | -2.18 | **0.03** | 0.90 |
| Age Group * Condition | -0.13 | 0.13 | -0.98 | 0.33 | 0.88 |
|  | Estimate | SE | z | p | Odds ratio |
| Model 2: Condition dummy-coded with reward as the reference group |  |  |  |  |  |
| Intercept | 0.65 | 0.09 | 6.90 | **<.01** | 1.91 |
| Curiosity (mean-centered) | 0.17 | 0.04 | 4.59 | **<.01** | 1.18 |
| Age group (effect-coded; YA: -1, OA: 1) | 0.03 | 0.09 | 0.32 | 0.75 | 1.03 |
| Condition (dummy-coded; Con: 1, Rew: 0) | -0.29 | 0.13 | -2.19 | **0.03** | 0.75 |
| Curiosity * Age Group | <0.01 | 0.03 | 0.02 | 0.98 | 1.00 |
| Curiosity * Condition | 0.11 | 0.05 | 2.18 | **0.03** | 1.11 |
| Age Group * Condition | 0.13 | 0.13 | 0.98 | 0.33 | 1.14 |

*Note:* YA = Younger adults; OA = Older adults; Con = Control; Rew = Reward

### Bayes Factor

To assess the relative evidence in support of the null or alternative hypothesis, we conducted a Bayes factor (BF) analysis on aggregated trivia recall and face recognition data. BF quantifies the relative likelihood of the observed data under the alternative hypothesis and the null hypothesis (Lakens et al., 2020). Therefore, BF > 1 indicates greater support for the alternative hypothesis, and BF < 1 indicates greater support for the null hypothesis. Following Jeffreys’ (1961) conventions, evidence for the alternative hypothesis can be qualitatively labeled as none (BF = 1), anecdotal (1 < BF ≤ 3), moderate (3 <BF ≤ 10), strong (10 <BF ≤ 30), very strong (30 <BF ≤ 100), or extreme (1 < BF ≤ 100). BF analyses were carried out in R version 1.3.959, using the BayesFactor package (Morey & Rouder, 2018) to calculate a JZS BF ANOVA with default prior scales (see also Galli et al., 2018).

### Trivia Recall

Results of the logistic mixed model estimation are reported in Tables 3 and 4. The final model regressed trial-level recall accuracy on trial-level curiosity rating, age group, and condition, as well as their two-way interaction terms. The three-way interaction term was not significant and was dropped to improve model AIC. Additionally, background measures that exhibited significant age-group differences were separately included as predictors in the model (see Table 1; education in years, verbal intelligence as measured by the Shipley vocabulary scale, processing speed as measured by correct responses on the Digit symbol substitution task, and affective state as measured by the Positive and Negative Affect Scale). None of the background measures impacted the pattern of results, and therefore results will be reported from the most parsimonious model without covariates. There were significant main effects of curiosity rating, Wald χ (1) = 80.38, *p* < 0.001, and condition, Wald χ (1) = 5.36, *p* = 0.02. These effects were qualified by a significant Curiosity x Condition interaction, Wald χ (1) = 4.74, *p* = 0.03. The effects of age group (see Figure 3b) and its two-way interactions were not significant, Age: Wald χ (1) = 2.04, *p* = 0.15; Age x Condition: Wald χ (1) = 0.95, *p* = 0.33; Age x Curiosity: Wald χ (1) < .01, *p* = 0.98.

**Table 3.** Parameter estimates for the final logistic mixed models predicting trial-level face recognition and trivia recall from curiosity, condition, and age group

|  | Estimate | SE | z | p | Odds ratio |
| --- | --- | --- | --- | --- | --- |
| Trial-level face recognition accuracy |  |  |  |  |  |
| Intercept | 0.29 | 0.08 | 3.52 | **<.01** | 1.34 |
| Curiosity (mean-centered) | 0.01 | 0.01 | 0.58 | 0.56 | 1.01 |
| Age group (effect-coded; YA: -1, OA: 1) | 0.10 | 0.08 | 1.22 | 0.22 | 1.11 |
| Condition (effect-coded; Con: -1, Rew: 1) | 0.07 | 0.08 | 0.94 | 0.35 | 1.08 |
| Curiosity * Age Group | -0.01 | 0.01 | -0.75 | 0.45 | 0.99 |
| Curiosity * Condition | -0.01 | 0.01 | -0.65 | 0.52 | 0.99 |
| Condition * Age Group | 0.03 | 0.08 | 0.41 | 0.68 | 1.03 |
|  | Estimate | SE | z | p | Odds ratio |
| Trial-level trivia recall accuracy |  |  |  |  |  |
| Intercept | 0.50 | 0.07 | 7.57 | **<.01** | 1.65 |
| Curiosity (mean-centered) | 0.22 | 0.03 | 8.66 | **<.01** | 1.25 |
| Age group (effects-coded; YA: -1, OA: 1) | 0.09 | 0.07 | 1.43 | 0.15 | 1.10 |
| Condition (effects-coded; Con: -1, Rew: 1) | 0.15 | 0.07 | 2.19 | **0.03** | 1.16 |
| Curiosity * Age Group | <0.01 | 0.03 | 0.02 | 0.98 | 1.00 |
| Curiosity * Condition | -0.05 | 0.02 | -2.18 | **0.03** | 0.95 |
| Age Group * Condition | -0.06 | 0.07 | -0.98 | 0.33 | 0.94 |

*Note:* YA = Younger adults; OA = Older adults; Con = Control; Rew = Reward

To probe the Curiosity x Condition interaction, two additional models were estimated with the control condition dummy-coded as 0 and the reward condition coded as 1, and vice versa, to isolate the curiosity effect within each condition. Odds ratios from these models indicated that a 1-unit increase in curiosity ratings increased the probability of correct recall by a factor of 1.31 in the control condition, but only by a factor of 1.18 in the reward condition. In other words, as curiosity increased, the odds that a trivia item was recalled increased more in the control condition than in the reward condition (Figure 4; see Table 4 for the estimates from the models used to probe the interaction). Consistent with our hypothesis, we replicated the interaction reported by Murayama and Kuhbandner (2011), but unexpectedly, the effect occurred in both age groups.

**Figure 4.**
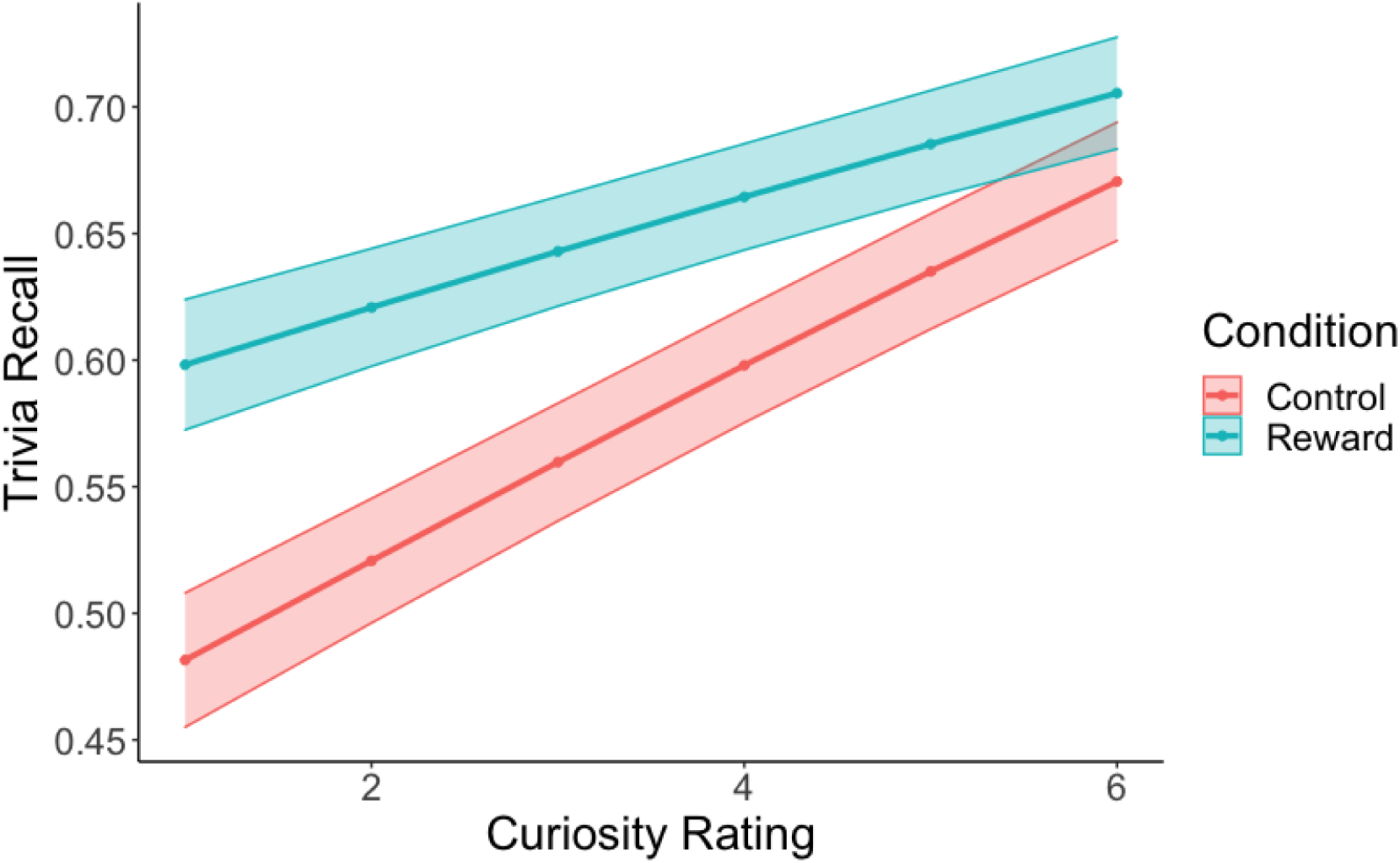
Predicted probability of correct trivia recall according to curiosity rating and condition. Compared to the control condition, the extrinsic reward condition produced a higher probability of correct recall of low-curiosity information, but this advantage disappeared for high-curiosity information. Confidence bands represent the standard error of the mean.

A JZS BF ANOVA revealed moderate evidence that the intercept model was preferred to the model that included the main effect of age (BF_01_ = 5.9), whereas extreme evidence supported the model that included the main effects of curiosity and condition (BF_10_ = 380.6). More importantly, the Curiosity x Condition interaction model was moderately preferred to the main effects model (BF_10_ = 3.8) and to the Curiosity x Condition x Age interaction model (BF_10_ = 3.5). Thus, we found moderate evidence against the hypothesis that curiosity and condition were differentially with associated with recall in younger and older adults.

### Face Recognition

Results of the logistic mixed model analysis for face recognition are reported in Table 3. The final model regressed trial-level accuracy (hits vs. misses) on trial-level curiosity rating, age group, and condition, as well as the two-way interaction terms. The three-way interaction term was non-significant and was dropped to reduce model complexity and improve model AIC. As in the trivia recall analysis, background measures with significant age-group differences (see Table 1) were separately included as predictors in the model. None of the background measures impacted the pattern of results, and therefore results are reported from the most parsimonious model without any covariates. Results from this model indicated no significant effects of curiosity rating, Wald χ (1) = 0.44, *p* = 0.51, age group, Wald χ (1) = 1.49, *p* = 0.22, or condition, Wald χ (1) = 0.83, *p* = 0.36. Likewise, none of the interactions were significant; Curiosity x Condition, Wald χ (1) = 0.42, *p* = 0.52; Curiosity x Age, Wald χ (1) = 0.56, *p* = 0.45; Condition x Age, Wald χ (1) = 0.17, *p* = 0.68. In other words, contrary to our hypotheses, an increase in the curiosity rating during encoding did not increase the odds of subsequent successful face recognition (odds ratio = 1.01).

The JZS BF ANOVA revealed extreme evidence for the intercept model to be preferred to the model that included the main effects of age, condition and curiosity (BF_01_ = 210.41), and the intercept model was extremely preferred to the interaction model (BF_01_ = 252.0). Thus, this provides extreme evidence against the hypothesis that recognition of irrelevant faces was affected by age, curiosity, condition or their interaction.

In summary, the likelihood of recognizing incidentally encoded faces was not modulated by differences in intrinsic and extrinsic motivation at encoding. By contrast, the likelihood of recalling trivia was greater for high-curiosity trivia, and for participants in the reward condition. However, the curiosity-related recall enhancement was stronger for participants in the control condition than for those in the reward condition. This pattern was similar in younger and older adults, a conclusion supported by the Bayes factor analysis.

## Discussion

The current study examined the effects of curiosity, an intrinsic motivator, and financial reward, an extrinsic motivator, on memory in younger and older adults. Participants encoded trivia that elicited varying levels of curiosity, and they later received a surprise trivia recall test. Half of the participants were also motivated extrinsically by the opportunity to earn money for correctly guessing answers to questions. Both curiosity and reward were associated with better trivia recall in both age groups, but the effects were interactive, such that reward enhanced memory for low-curiosity trivia, but not for high-curiosity trivia. The hypothesis of a reduced influence of reward on memory, in older adults as compared to younger adults, was not confirmed. Finally, memory for unrelated faces encountered during the study phase was not modulated by curiosity or reward, in either age group.

There are several possible reasons why we did not find an association between curiosity and incidental face memory. First, past studies using similar tasks (Galli et al., 2018, Exp. 1; Gruber et al., 2014; Murphy et al., 2020; Stare et al., 2018) have yielded small effects of curiosity on memory for unrelated information (η_p_^2^ ≤ .08). In light of the relatively small sample sizes in the current study and in past published studies (*n* ≤ 40 per age group), null findings are therefore not unexpected. For example, the only prior study to report a curiosity effect on memory for unrelated faces in younger and older adults failed to replicate this effect in a second experiment (Galli et al., 2018).

There were also several methodological differences between the current study and previous published studies. First, our task borrowed aspects of a procedure used by Murayama and Kuhbandner (2011) to test the interaction of curiosity and financial reward on memory.

Therefore, the incidental face stimulus and the trivia answer were separated by a guessing task (“Do they know the answer?”). If the effect of curiosity on incidental face memory depends more on curiosity satisfaction (Marvin & Shohamy, 2016), surprise (Baranes et al., 2015), or post-answer interest (McGillivray et al., 2015) than on the anticipatory affect associated with curiosity, then the guessing task may have disrupted the effect. It should be noted, however, that one recent study (Murphy et al., 2020) suggests that incidental memory enhancement from curiosity is contingent on proximity to curiosity elicitation rather than curiosity satisfaction.

Another difference between the current paradigm and prior studies was that the experimenter acted as a scribe. Rather than interfacing with the task directly, participants dictated their responses aloud to the experimenter to enter on their behalf. This procedure may have influenced participants’ level of engagement with face stimuli and introduced an element of social desirability when participants had to indicate whether the person shown in the picture was likely to know the answer to the trivia question (e.g., “I don’t want to seem judgmental, so I’ll say yes”).

It is also possible that we were unable to detect an effect of our manipulations because of the low performance on the face recognition task (corrected recognition *M* = 46.2%). However, these data represent better performance than prior studies that identified an effect of curiosity on irrelevant face recognition (Gruber et al., 2014, *M =* 40.1%; Galli et al., 2018, Exp 1, *M =* 31.5%) making it unlikely that low performance was a large factor.

An alternative perspective on the current findings relates to memory selectivity effects of reward anticipation and arousal. According to ABC theory, noradrenergic arousal, associated with states of reward anticipation (Knutson & Greer, 2008) and curiosity (Sakaki et al., 2018) biases attention and memory processes toward task-relevant stimuli while suppressing task-irrelevant stimuli (Mather & Sutherland, 2011). However, this perspective is not consistent with our results, as it would predict greater memory selectivity associated with high-curiosity than low-curiosity trials. In other words, according to ABC theory, we would expect better recognition for faces encountered in low-curiosity trials compared to high-curiosity trials during which attention and memory would presumably be more selective.

A final possibility relates to the confound between task relevance (trivia vs. faces) and test format (recall vs. recognition). It is possible that recall is more susceptible to motivational influences than recognition. To eliminate the confound between task relevance and test format, future studies could manipulate task relevance while holding test format constant, and vice versa. It should be noted, however, that past studies reporting curiosity-related enhancement of incidental face memory also used recognition tests (Galli et al., 2018, Exp. 1; Gruber et al., 2014; Murphy et al., 2020; Stare et al., 2018). Moreover, incentive effects on incidental recognition memory tests have been documented in younger and older adults (Mather & Schoeke, 2011). In sum, of the potential explanations for the null effect of curiosity on incidental face memory in the current study, low statistical power to detect small effects is the most likely candidate. Replication with a larger sample, and meta-analytic synthesis across published studies, are the most promising avenues for resolving the question of whether, and to what extent, curiosity benefits incidental learning of unrelated material in younger and older adults.

Which mechanisms may account for the interaction of curiosity and reward in younger and older adults’ trivia recall? One interpretation is that monetary incentives are less effective in tasks with high intrinsic value, compared to tasks with low intrinsic value (e.g., Murayama & Kuhbandner, 2011). Inherent in this view is the notion of a trade-off between the effects of intrinsic and extrinsic motivation on cognitive performance, as predicted by self-determination theory (for a review, see Deci et al., 1999). However, the idea that monetary incentives “crowd out” the benefits of curiosity has not been unchallenged. One recent study with younger adults reported additive effects of curiosity and reward on trivia recall (Duan et al., 2020), suggesting that the two sources of motivation can work in tandem. In Duan et al.’s (2020) study, encoding was intentional, reward was manipulated within subjects and it was contingent on correct recall in the test phase. In contrast, in our study and in the study by Murayama and Kuhbandner (2011), encoding was incidental, reward was manipulated between subjects, and its delivery was contingent on correct guesses during the study phase. First, intentional encoding success may depend more strongly on strategic processes than incidental encoding. Second, since curiosity operates during encoding, providing the performance-contingent bonus at retrieval rather than encoding may reduce the chance of interaction between reward and curiosity. Thus, it is possible that the interaction occurs specifically under incidental learning conditions, and only when intrinsic/extrinsic motivators overlap (both at encoding, rather than one at encoding and one at retrieval). How differences in study and test format influence the interaction of intrinsic and extrinsic motivation on learning and memory is an important question for future work.

Regardless of the precise nature of the interaction between reward and curiosity, our findings suggest that intrinsic motivation to learn from curiosity can produce an almost identical increase in the probability of correct recall as extrinsic motivation from reward. Therefore, interventions aimed at enhancing memory and learning outcomes can target intrinsic motivational states (e.g., curiosity, interest, satisfaction, and surprise; Ozono, et al., 2020; Ryan & Deci, 2000) instead of extrinsic contingencies of reinforcement (e.g., monetary bonuses, testing grades, social recognition; Slavin, 2010; Stan, 2012).

Lastly, the interaction of curiosity and reward on memory was similar in younger and older adults, in contradiction of our hypothesis that older adults’ memory would not be influenced by the provision of financial incentives. Although statistical null effects of age do not necessarily indicate the absence of age differences (Lakens et al., 2020), the Bayes factor analyses lent support to the models without main effects of age or its higher-order interactions. This finding aligns with prior reports of age-related similarities in the effects of intrinsic and extrinsic motivation on memory (Galli et al., 2018, Exp 2; Mather & Schoeke, 2011; McGillivray et al., 2015; Spaniol et al., 2014). These observations are particularly interesting in the face of well-documented dopaminergic decline and changes in reward-related brain activation (Bäckman et al., 2010; Dreher et al., 2008; Eppinger et al., 2011), age differences in temporal discounting of reward (Eppinger et al, 2012), and changes in motivational priorities across the lifespan (Carstensen et al., 1999), which may increase older adults’ preference for non-financial rewards (e.g., social rewards; Rademacher et al., 2014). The current findings suggest that, despite age-related reductions in reward-related processing, both intrinsic curiosity and financial reward are effective methods for boosting memory in older adults.

The study had several limitations. First, its sample size, while based on an a priori power analysis and comparable to other studies, was relatively small. Although the study was sufficiently powered to detect the two-way interaction of interest within each age group, a larger sample size would have strengthened our conclusion about the three-way interaction of age, curiosity, and reward condition. Second, the findings may not generalize beyond the populations included in this research: predominantly White, urban, highly educated, and healthy younger and older adults. Third, the decision to use the experimenter as a scribe in the incidental encoding task may have differentially influenced participants’ level of engagement with the trivia and face stimuli to a degree that we are unable to measure and account for in the current design. Last, the study used behavioural measures only. In future work, a multi-method approach to studying the impact of extrinsic reward on curiosity-enhanced memory could shed light on the neural substrates of the two types of motivational influences, which are likely overlapping (but see Duan et al., 2020). Neuroimaging would help characterize the interaction of intrinsic curiosity and extrinsic reward in the brain, and could pinpoint age differences in the neural substrates of interactions between curiosity, reward, and memory (Sakaki et al., 2018). Similarly, pupillometry and eye-tracking could shed light on the contributions of curiosity-related arousal and attention to subsequent memory performance (Baranes et al. 2015; Kang et al., 2009). For example, different mechanisms may be driving curiosity-driven memory enhancement of task-relevant and irrelevant information. If this were the case, one prediction might be that pupil dilation associated with pre-answer anticipation predicts trivia recall, whereas dilation associated with post-answer surprise/satisfaction predicts face recognition performance. However, this pattern may manifest differently in older adults who undergo changes in the locus-coeruleus norepinephrine system responsible for arousal (Lee et al., 2018). In sum, insight into age differences in the mechanisms giving rise to curiosity-memory interactions will benefit from a combination of behavioral and physiological measures.

To conclude, the current study demonstrates that curiosity is a powerful predictor of incidental memory in younger and older adults, but that the curiosity benefit may be narrow rather than broad. This finding holds implications for educational practice with adult learners. For example, rather than interspersing lectures with curiosity-arousing tidbits to improve learners’ memory for dry material, a more effective strategy may be to arouse curiosity about the material itself. Another key implication derives from the interaction of curiosity and reward effects on memory, which was present in both age groups. While the prospect of financial reward improved memory performance, it did so only for uninteresting material. In the context of adult learning, this suggests that extrinsic incentives should be used only as a last resort (i.e., when it is difficult or impractical to make the curriculum interesting or self-relevant to learners). Overall, these findings complement and extend knowledge about motivation-cognition interactions in healthy aging, and they highlight the importance of curiosity for cognitive function across the adult lifespan (Sakaki et al., 2018).

## Appendix A

List of trivia items used:

| Question | Answer | Trivia Domain | Answer Type | Mean Curiosity Rating |  |  |  |
| --- | --- | --- | --- | --- | --- | --- | --- |
|  |  |  |  | Younger |  | Older |  |
|  |  |  |  | M | SD | M | SD |
| 'Arco irisa' is the Spanish term for what natural phenomenon? | Rainbow | General | Categorical | 4.05 | 1.54 | 3 | 1.73 |
| 'Circle of Life' came from which film? | Lion King | Pop Culture | Categorical | 4.13 | 1.89 | 2.6 | 1.3 |
| 'Come as You Are', a song by the grunge band Nirvana was released on which album? | Nevermind | Pop Culture | Categorical | 2.27 | 1.91 | 2.06 | 1.21 |
| 'My heart will go on' came from which movie? | Titanic | Pop Culture | Categorical | 2.17 | 1.6 | 2 | 1.41 |
| 'The Fresh Prince of' where was the subject of a sitcom of 140+ shows? | Bel-Air | Pop Culture | Categorical | 3.67 | 1.86 | 2.2 | 0.84 |
| A couple celebrating their crystal wedding anniversary have been married for how many years? | 15 | General | Numeric | 3.17 | 1.72 | 3.26 | 1.79 |
| A goat sucker is what type of animal? | A bird | Animals | Categorical | 4 | 1.64 | 3.76 | 1.73 |
| A passive two-terminal electrical component that produces electrical resistance in a circuit is called what? | A resistor | General | Categorical | 3.88 | 1.81 | 3.23 | 1.9 |
| A phlebotomist extracts what from the human body? | Blood | STEM | Categorical | 3.36 | 1.75 | 4.43 | 1.65 |
| A Shakespearean sonnet consists of how many lines? | 14 | General | Numeric | 3.53 | 1.88 | 3.53 | 1.84 |
| A typical mayfly lives for how many days? | One | General | Numeric | 3.57 | 1.4 | 2.27 | 0.79 |
| Al Gore was elected Senator for what state in 1985? | Tennessee | History | Categorical | 3.03 | 1.68 | 3 | 1.8 |
| Andrew Carnegie developed the iron and steel industry in which city? | Pittsburgh | History | Categorical | 3.22 | 1.86 | 4.29 | 1.38 |
| At what age is it mandatory to retire from the Supreme Court of Canada? | 75 | General | Numeric | 3.17 | 1.76 | 3.67 | 1.21 |
| Bogota is the high altitude capital of which country? | Colombia | Geography | Categorical | 3 | 1.48 | 2 | 0 |
| By law, what is banned in Japanese restaurants? | Tipping | General | Categorical | 5.24 | 1.14 |  |  |
| Dendrophobia is the fear of what? | Trees | General | Categorical | 4.55 | 1.64 | 4.67 | 1.35 |
| Emerald is the birthstone for which month? | May | General | Categorical | 3.65 | 1.75 | 2.44 | 1.41 |
| Father's Day was first celebrated in which country? | USA | History | Categorical | 4.11 | 1.45 | 2.85 | 1.82 |
| Giorgio Armani trained for which profession although he didn't qualify? | Doctor | History | Categorical | 3.18 | 1.94 | 2.92 | 1.66 |
| Granadilla is another name for which fruit? | Passionfruit | General | Categorical | 4.14 | 1.55 | 4.07 | 1.59 |
| Hale-Bopp is classified as which type of small solar system body? | Comet | STEM | Categorical | 3.63 | 1.77 | 3.14 | 1.7 |
| How are the Motion Picture Academy Awards also known? | Oscars | Pop Culture | Categorical | 2.8 | 1.62 | 5 |  |
| How many dots are there on two dice? | 42 | General | Numeric | 1 | 1.02 | 3 | 1.58 |
| How many friends are there in the sitcom "Friends"? | 6 | Pop Culture | Numeric | 3.2 | 1.3 | 2.13 | 1.19 |
| How many letters are there in the German alphabet? | 30 | General | Numeric | 2.88 | 1.73 | 3.9 | 1.62 |
| How many points does the maple leaf on the Canadian flag have? | 11 | General | Numeric | 2.9 | 1.29 | 4.57 | 1.62 |
| How many red stripes are there on the United States flag? | 7 | General | Numeric | 3.79 | 1.76 | 2.86 | 1.56 |
| How many squares are on a standard chessboard? | 64 | General | Numeric | 3.73 | 1.56 | 3.43 | 1.79 |
| How many symphonies did Beethoven compose? | 9 | General | Numeric | 4.09 | 1.76 | 3.54 | 1.56 |
| How many times zones does France and its overseas territories use? | 12 | Geography | Numeric | 3.22 | 1.93 | 2.8 | 1.79 |
| How many U.S. states border the Gulf of Mexico? | 5 | Geography | Numeric | 2.55 | 1.34 | 3.44 | 1.59 |
| How many years were there between the two Los Angeles Olympics? | 52 | Sports | Numeric | 2.86 | 1.46 | 2.33 | 1.33 |
| How may dots are there on two dice? | 42 | General | Numeric | 2.5 | 1.29 |  |  |
| Ice hockey pucks are made from what material? | Vulcanized rubber | Sports | Categorical | 3.11 | 2.15 | 4.71 | 0.49 |
| In 2008, which country hosted a Formula 1 Race for the first time? | Singapore | Sports | Categorical | 2.89 | 1.77 | 2.4 | 1.35 |
| In baseball, where do the Blue Jays come from? | Toronto | Sports | Categorical | 4 | 1.41 |  |  |
| In boxing, what is the term for an illegal punch to the back of the head or base of the skull? | Rabbit Punch | Sports | Categorical | 3.1 | 1.62 | 3.1 | 1.37 |
| In British mythology, who were Gog and Magog? | Giants | General | Categorical | 3.35 | 2.06 | 2.72 | 1.78 |
| In economics, what does the letter F stand for in IMF? | Fund | Business | Categorical | 2.5 | 1.68 | 3.75 | 1.5 |
| In fiction, what is the last name of Dr. Hannibal--the Cannibal? | Lecter | Pop Culture | Categorical | 2.17 | 1.34 | 2.33 | 1.66 |
| In football, what position is QB? | Quarterback | Sports | Categorical | 6 |  | 3 |  |
| In our solar system, which planet has the shortest day? | Jupiter | STEM | Categorical | 4.87 | 1.36 | 4.05 | 1.81 |
| In square miles, how big is Lake Ontario? | 7550 | STEM | Numeric | 2.86 | 1.68 | 3.25 | 1.86 |
| In the 1700's, what did some residents of London purchase to avoid going to hell? | Insurance | History | Categorical | 4.1 | 1.68 | 4.19 | 1.6 |
| In the classic board game Monopoly, how much does it cost to buy a railroad? | 200 | General | Numeric | 1.8 | 1.19 | 2.63 | 1.54 |
| In the human body what is the hallux? | Big toe | STEM | Categorical | 3.85 | 1.93 | 4.36 | 1.5 |
| In the United States and Canada, one ton is a unit of measure that contains how many pounds? | 2000 | STEM | Numeric | 3.8 | 1.74 | 3.6 | 2.3 |
| In what country did Gordon Ramsay open his first restaurant in 2007? | Ireland | General | Categorical | 3.69 | 1.82 | 2.6 | 1.3 |
| In what country was the first written account of children using the phrase 'trick or treat' on Halloween? | Canada | General | Categorical | 3.93 | 1.91 | 3.67 | 1.32 |
| In what country were checkers invented? | Egypt | General | Categorical | 3.68 | 1.81 | 4 | 1.46 |
| In what country would you find large ancient geoglyphs known as the Nazca Lines? | Peru | Geography | Categorical | 3.82 | 1.94 | 4.25 | 1.66 |
| In what sport does a jammer score a point for each opponent she skates past? | Roller Derby | Sports | Categorical | 3.5 | 1.96 | 2.8 | 1.42 |
| In what year was Instagram launched? | 2010 | Pop Culture | Numeric | 2.95 | 1.58 | 2.75 | 1.77 |
| In what year was Stephen Harper first elected as Prime Minister? | 2006 | General | Numeric | 2.88 | 1.88 | 2.95 | 1.64 |
| In which branch of the arts is Katherine Dunham famous? | Ballet | Pop Culture | Categorical | 2.67 | 1.49 | 3.19 | 1.69 |
| In which country did cheddar cheese originate? | England | History | Categorical | 3.82 | 1.78 | 4.25 | 0.96 |
| In which country is an Afghani a unit of currency? | Afghanistan | General | Categorical | 3.5 | 3.54 | 2.33 | 2.31 |
| In which country is the Simpson Desert found? | Australia | Geography | Categorical | 3 | 1.69 | 3.12 | 1.96 |
| In which month of the year is Battle of Britain week? | September | General | Categorical | 3.04 | 1.74 | 3.13 | 1.78 |
| In which season do most burglaries take place? | Winter | General | Categorical | 5.1 | 1.29 | 4.5 | 1.32 |
| In which Tennessee city is it illegal to lasso fish? | Knoxville | Geography | Categorical | 3.16 | 1.94 | 2.52 | 1.81 |
| In which US state is John F Kennedy buried? | Virginia | General | Categorical | 3.59 | 1.94 | 3.5 | 1.58 |
| Jack the Ripper is the name given to an unidentified serial killer that terrorized what city in 1888? | London, England | History | Categorical | 3.6 | 1.96 | 1 | 0 |
| Karakul, Texel and Romney Marsh are different kinds of what? | Sheep | Animals | Categorical | 3.18 | 1.94 | 2.81 | 1.72 |
| Lagnoperissia is a fancy name for what sexual condition? | Nymphomania | STEM | Categorical | 4.3 | 1.59 | 3 | 1.66 |
| Marxist revolutionary Che Guevara was born in what country? | Argentina | History | Categorical | 4 | 1.92 | 4.23 | 1.59 |
| Most electrogenic fish are also what? | Electroreceptive | STEM | Categorical | 3.96 | 1.82 | 3.87 | 1.41 |
| Nariyal is the Indian term for which nut? | Coconut | General | Categorical | 3.84 | 1.68 | 3.11 | 1.32 |
| Olympia is the capital city of which U.S. state? | Washington | Geography | Categorical | 4.19 | 1.42 | 3.25 | 2.12 |
| On a computer keyboard, what letter is between Q and E? |  |  | Categorical |  |  | 3.4 | 1.67 |
| On average, how much time will an American spend stopped at a red light over their whole life? | 2 weeks | General | Numeric | 3.61 | 2.01 | 2.85 | 1.83 |
| On Sunday, in Columbus Ohio, it is illegal to sell what? | Cornflakes | General | Categorical | 4.34 | 1.43 | 3.21 | 1.65 |
| On the Apollo 11 moon mission, which astronaut stayed aloft in the command module while Neil Armstrong and Buzz Aldrin walked on the moon? | Michael Collins | History | Categorical | 3.91 | 1.82 | 3.86 | 1.56 |
| On what day of the week do most automobile accidents happen? | Saturday | General | Categorical | 5.11 | 1.05 | 4.61 | 1.58 |
| Papua New Guinea is bordered by which country to the west? | Indonesia | Geography | Categorical | 3.62 | 1.7 | 3.67 | 1.75 |
| Ranch dressing originated in which U.S. state? | Alaska | History | Categorical | 3.38 | 1.96 | 2.54 | 1.35 |
| Robert Gallo was one of the pioneers in the identification of which virus? | HIV | History | Categorical | 3.38 | 1.75 | 3.77 | 1.54 |
| Sika, fallow, and Roe, are what types of animal? | Deer | Animals | Categorical | 3.88 | 1.45 | 4.57 | 1.4 |
| Someone who suffers from oneirophobia is scared of what? | Dreams | General | Categorical | 4.26 | 1.76 | 3.67 | 1.84 |
| Sriracha is type of hot sauce named after a city located in what country? | Thailand | General | Categorical | 4.88 | 1.22 | 3.94 | 1.35 |
| The aardvark is native to which continent? | Africa | Geography | Categorical | 4.38 | 1.72 | 4.83 | 1.59 |
| The American musician Little Walter was associated with which instrument? | Harmonica | History | Categorical | 2.53 | 1.25 | 3.63 | 1.54 |
| The chemist Alfred Nobel made his money by inventing what? | Dynamite | STEM | Categorical | 3.6 | 1.72 | 4.89 | 0.6 |
| The expression 'Oy vey!' comes from which language? | Yiddish | General | Categorical | 4.15 | 1.6 | 4.25 | 0.96 |
| The famous American writer Samuel Langhorne Clemens is better known by what pen name? | Mark Twain | Pop Culture | Categorical | 3 | 1.66 | 3.88 | 1.25 |
| The first film star to appear on a postage stamp was who? | Grace Kelly | History | Categorical | 3.09 | 1.73 | 3.32 | 1.53 |
| The Gobi Desert is primarily situated in which country? | Mongolia | Geography | Categorical | 3.82 | 1.59 | 5.31 | 0.48 |
| The Hound of the Baskervilles is a crime novel featuring which fictional detective? | Sherlock Holmes | Pop Culture | Categorical | 3 | 1.69 | 4 | 0 |
| The May Queen, Wisley Crab, Lane's Prince Albert and Foxwhelps are all species of what? | Apple | General | Categorical | 2.79 | 1.65 | 4.95 | 1.23 |
| The Mediterranean Sea connects to the Red Sea through which waterway? | Suez Canal | Geography | Categorical | 4.04 | 1.72 | 4.14 | 1.86 |
| The men's magazine GQ was formerly known by what longer name? | Gentlemen's Quarterly | Pop Culture | Categorical | 2.96 | 2.07 | 2.38 | 1.19 |
| The most Asian elephants to be found in their natural habitat can be found in what country? | India | Geography | Categorical | 3.67 | 1.91 | 5.3 | 1.57 |
| The Principality of Monaco is a sovereign city-state bordered on three sides by which country? | France | Geography | Categorical | 3.19 | 1.8 | 4.2 | 1.48 |
| The sale of what was prohibited in America during prohibition? | Alcohol | History | Categorical | 4 | 2 |  |  |
| The study of birds eggs is called what? | Oology | Animals | Categorical | 3.76 | 1.73 | 3.78 | 1.59 |
| The US declared war on which country after the bombing of Pearl Harbor? | Japan | History | Categorical | 3.33 | 2.52 |  |  |
| The world's fastest growing plant is a species of what? | Bamboo | General | Categorical | 4.17 | 1.38 | 4.69 | 1.66 |
| Unleavened bread or matzo, used for the Jewish feast of Passover, does not include what ingredient? | Yeast | Religion | Categorical | 3 | 1.83 | 1.75 | 0.96 |
| Unleavened bread or matzo, used for the Jewish feast of Passover, does not include what? | Yeast | Religion | Categorical | 4 | 1.22 |  |  |
| Walt Disney's famous deer was named what? | Bambi | Pop Culture | Categorical | 1.8 | 1.3 | 4 | 1.41 |
| What 20th-century conflict was dubbed the "forgotten war" despite 54,246 U.S. deaths? | The Korean War | History | Categorical | 3.63 | 1.86 | 4.5 | 2.07 |
| What are the small indentations on a golf ball called? | Dimples | Sports | Categorical | 3.56 | 1.92 | 3.29 | 1.57 |
| What brave-hearted Scottish patriot led soldiers to a defeat of the English at the Battle of Cambuskenneth in 1297? | William Wallace | History | Categorical | 3.04 | 1.82 | 3.23 | 1.59 |
| What breed of horse is best known for its use in racing? | Thoroughbred | Sports | Categorical | 2.64 | 1.62 | 4.22 | 1.3 |
| What California city did the last Pony Express ride end in? | Sacramento | History | Categorical | 2.26 | 1.48 | 2.89 | 1.59 |
| What came after "The Brady" in the sitcom title? | Bunch | Pop Culture | Categorical | 2.07 | 1.49 | 2.5 | 0.71 |
| What chemical element gives the blood of a lobster a bluish tint? | Copper | STEM | Categorical | 3.74 | 1.63 | 3.95 | 1.68 |
| What city had the first public school, college and newspaper in the thirteen British colonies? | Boston | History | Categorical | 2.75 | 1.45 | 4.2 | 1.26 |
| What colour are the stars on the United States of America flag? | White | General | Categorical | 3.33 | 1.53 | 3.43 | 2.23 |
| What colour is a giraffe's tongue? | Black | Animals | Categorical | 3.17 | 1.64 | 3.85 | 1.77 |
| What colour is the M in McDonald's logo? | Yellow | Pop Culture | Categorical | 3.5 | 2.12 | 2.22 | 1.3 |
| What continent is subjected to the world's largest ozone hole? | Antarctica | Geography | Categorical | 4.09 | 1.58 | 5 | 1.25 |
| What country gave Florida to the USA in 1891? | Spain | History | Categorical | 4 | 1.54 | 2 | 1.41 |
| What country was formerly known as Abyssinia? | Ethiopia | History | Categorical | 3.84 | 1.6 | 4.38 | 1.41 |
| What day in July is Canada Day? | July 1st | General | Numeric | 4.5 | 1.29 |  |  |
| What did Joseph Priestley discover in 1774? | Oxygen | History | Categorical | 3.25 | 1.98 | 3.75 | 1.44 |
| What did people in the middle ages throw at the bride and groom? | Eggs | History | Categorical | 4.52 | 1.69 | 3.91 | 1.44 |
| What did teenager Anne Frank leave behind that was published after her death? | Diary | History | Categorical | 2.33 | 1.15 | 2 |  |
| What diet drink was hyped by Coca-Cola for having only one calorie, in 1963? | Tab | History | Categorical | 3.7 | 1.45 | 3.71 | 1.53 |
| What do doctors look at through an ophthalmoscope? | The eye | General | Categorical | 4.11 | 1.56 | 5 | 1.41 |
| What do you call an angle more than 90 degrees and less than 180 degrees? | Obtuse | STEM | Categorical | 3.33 | 2.16 | 3.43 | 1.27 |
| What do you call the smell which wine gives off? | Bouquet | General | Categorical | 3.95 | 1.65 | 5 | 1.49 |
| What does a philatelist collect? | Stamps | General | Categorical | 4.14 | 1.65 | 1 |  |
| What does the C stand for in LCD? | Crystal | General | Categorical | 3.31 | 1.96 | 3 | 1.65 |
| What element begins with the letter "K"? | Krypton | STEM | Categorical | 4.15 | 1.34 | 4 | 1.87 |
| What explorer introduced Italians to spaghetti in the 14th century? | Marco Polo | History | Categorical | 3.75 | 1.73 | 3.88 | 1.55 |
| What explorer introduced pigs to North America? | Christopher Columbus | History | Categorical | 3.13 | 1.54 | 3.89 | 1.97 |
| What F-word is defined in physics as a 'nuclear reaction in which nuclei combine to form more massive nuclei'? | Fusion | STEM | Categorical | 4.67 | 1.73 | 3.25 | 1.26 |
| What flavour is cointreau? | Orange | General | Categorical | 3.56 | 1.92 | 3.88 | 1.46 |
| What Greek mathematician is considered the founder and father of Geometry? | Euclid | STEM | Categorical | 3.38 | 1.63 | 4.36 | 1.36 |
| What group of Pacific islands did Japan attack the day after Pearl Harbor? | The Philippines | History | Categorical | 3.62 | 1.33 | 4.36 | 1.63 |
| What holiday, celebrated December 26 to January 1, is named after the Swahili word for 'first'? | Kwanzaa | Religion | Categorical | 4.63 | 1.38 | 4.71 | 1.6 |
| What insect accurately indicates the air temperature? | Cricket | General | Categorical | 4.16 | 1.68 | 4.58 | 1.31 |
| What is the common term for the tennis ailment "lateral humeral epicondylitis"? | Tennis Elbow | Sports | Categorical | 3.35 | 1.84 | 2.86 | 2.04 |
| What is a group of jellyfish called? | A smuck | Animals | Categorical | 4 | 1.37 | 3.73 | 1.86 |
| What is actor Woody Harrelson's middle name? | Tracy | Pop Culture | Categorical | 2.84 | 1.86 | 1.91 | 1.44 |
| What is Bono's real name? | Paul Hewson | General | Categorical | 3.07 | 1.82 | 2.96 | 1.73 |
| What is it called when a player scores two goals in a game of soccer? | A brace | Sports | Categorical | 3.08 | 1.79 | 3.3 | 1.49 |
| What is rum distilled from? | Sugar cane | General | Categorical | 4.75 | 1.65 | 3.7 | 1.25 |
| What is someone who shoes horses called? | A farrier | General | Categorical | 3.74 | 1.66 | 4 | 1.91 |
| What is the Aloha State? | Hawaii | Geography | Categorical | 3.83 | 2.14 |  |  |
| What is the capital city of Canada's Yukon territory? | Whitehorse | Geography | Categorical | 3.83 | 1.75 | 4 | 1.53 |
| What is the capital of Canada? | Ottawa | Geography | Categorical | 3 |  |  |  |
| What is the capital of Saskatchewan? | Regina | Geography | Categorical | 3.91 | 1.87 | 4.75 | 1.5 |
| What is the English translation for the name of the German automaker Volkswagen? | People's car | General | Categorical | 4 | 1.58 | 4.17 | 1.83 |
| What is the fastest animal in the world? | Peregrine Falcon | Animals | Categorical | 4.68 | 1.38 | 5.17 | 0.72 |
| What is the Great Barrier Reef made from? | Coral | Animals | Categorical | 3.4 | 2.41 |  |  |
| What is the heaviest naturally occurring element found on Earth? | Uranium | STEM | Categorical | 4.83 | 1.42 | 4 | 1.57 |
| What is the largest city in Alabama? | Birmingham | Geography | Categorical | 2.56 | 1.5 | 2.67 | 1.43 |
| What is the largest compartment in a cow's stomach called? | Rumen | Animals | Categorical | 3.85 | 1.46 | 3.33 | 1.61 |
| What is the largest compartment in a cow's stomach? | Rumen | Animals | Categorical | 4.43 | 1.51 |  |  |
| What is the largest country in Central America? | Nicaragua | Geography | Categorical | 4.14 | 1.66 | 4.2 | 1.48 |
| What is the largest freshwater lake in the world? | Lake Superior | Geography | Categorical | 4.33 | 1.4 | 4.67 | 1 |
| What is the largest sea in the world? | Philippine Sea | Geography | Categorical | 4.27 | 1.42 | 5.06 | 1.12 |
| What is the least popular month for U.S. weddings? | January | General | Categorical | 3.22 | 1.86 | 2.75 | 1.44 |
| What is the main colour on the Chinese flag? |  |  | Categorical |  |  | 1.5 | 0.71 |
| What is the most common non-contagious disease in the world? | Tooth decay | STEM | Categorical | 4.53 | 1.78 | 4.79 | 1.67 |
| What is the most popular sport in Canada? | Hockey | Sports | Categorical | 2 | 1.15 | 4 | 1.41 |
| What is the most purchased grocery item in Canada? | Kraft Dinner | General | Categorical | 5.14 | 1.39 |  |  |
| What is the most purchased non-perishable food item in Canada? | Ketchup | General | Categorical | 3 |  |  |  |
| What is the most purchased non-perishable pantry item in Canada? | Kraft Dinner | General | Categorical | 4.29 | 1.8 | 4.38 | 1.2 |
| What is the most widely eaten fish in the world? | Herring | General | Categorical | 4.33 | 1.43 | 5.75 | 0.45 |
| What is the name for a collection of frogs? | Army | Animals | Categorical | 3.52 | 1.78 | 4.19 | 1.7 |
| What is the name for Canadian bacon? | Peameal | General | Categorical | 3.9 | 1.71 | 3 | 1.51 |
| What is the name for meteoroids that survive entry through the atmosphere and reach Earth's surface? | Meteorites | STEM | Categorical | 3.85 | 1.66 | 4.24 | 1.71 |
| What is the name for the dish made with french fries, gravy, and cheese curds? | Poutine | General | Categorical | 1.32 | 2.05 | 1 | 3.24 |
| What is the name for the most popular Canadian coffee chain? | Tim Horton's | Animals | Categorical | 5 |  | 2 |  |
| What is the name for the offspring of a male donkey and a female horse? | Mule | Animals | Categorical | 4.72 | 1.49 | 3.62 | 1.33 |
| What is the name for the upper arm bone found in humans? | Humerus | STEM | Categorical | 3.82 | 1.51 | 4.73 | 1.01 |
| What is the name of the dog from the 1960s television cartoon The Jetsons? | Astro | Pop Culture | Categorical | 2.76 | 1.62 | 2.96 | 1.77 |
| What is the name used for the study of earthquakes? | Seismology | STEM | Categorical | 3.7 | 1.61 | 4.17 | 1.64 |
| What is the national animal of Canada? | Beaver | Geography | Categorical | 5.25 | 0.96 | 5 |  |
| What is the NHL? | National Hockey League | Sports | Categorical | 5 | 1.41 |  |  |
| What is the Northern Hemisphere equivalent to the Tropic of Capricorn? | Tropic of Cancer | General | Categorical | 3.05 | 1.85 | 2.5 | 1.29 |
| What is the official language of Greenland? | Greenlandic | General | Categorical | 4.43 | 1.41 | 4.5 | 1.4 |
| What is the oldest known tree food? | Walnut | General | Categorical | 3.67 | 1.8 | 4 | 1.52 |
| What is the only bird known to fly backwards? | Hummingbird | Animals | Categorical | 4.85 | 1.86 | 4.78 | 1.09 |
| What is the only word in English ending in the letters 'mt'? | Dreamt | General | Categorical | 4 | 1.91 | 5.33 | 1.24 |
| What is the plastic sheath at the end of a shoelace called? | An Aglet | General | Categorical | 4.45 | 1.85 | 4 | 1.62 |
| What is the postal abbreviation for California? | CA | General | Categorical | 3.09 | 1.97 | 2.67 | 1.03 |
| What is the shallowest ocean in the world? | Arctic Ocean | Geography | Categorical | 3.61 | 1.79 | 4.53 | 1.65 |
| What is the slang name for a 1 dollar Canadian coin? | Loonie | General | Categorical | 4 | 2.83 |  |  |
| What is the tallest free-standing structure in Canada? | CN Tower | General | Categorical | 3.5 | 0.71 |  |  |
| What is the wobbly red piece of flesh under the beak of a turkey? | A wattle | Animals | Categorical | 4 | 1.8 | 4.11 | 1.54 |
| What Italian astronomer invented the thermometer in 1592? | Galileo | History | Categorical | 3.93 | 2.09 | 4.44 | 1.51 |
| What item is banned only during Halloween from 12am October 31st to 12pm November 1st in Hollywood California? | Silly string | General | Categorical | 4.64 | 1.5 | 3.73 | 1.67 |
| What kind of animal is the emblem of the US democratic political party? | Donkey | General | Categorical | 2.64 | 1.57 | 3.21 | 1.67 |
| What kind of codes did American supermarkets introduce in the mid 70s? | Barcodes | General | Categorical | 2.8 | 1.87 | 1.67 | 1.15 |
| What kind of leaf is on the Canadian flag? | Maple | General | Categorical | 4.5 | 0.71 |  |  |
| What kind of plant bulbs were once exchanged as a form of currency? | Tulips | History | Categorical | 4.67 | 1.44 | 4.5 | 1.87 |
| What kind of plant does the Colorado beetle attack? | Potato | General | Categorical | 3.11 | 1.83 | 2.9 | 1.55 |
| What measure of energy comes from the Latin word meaning 'heat'? | The calorie | General | Categorical | 3.96 | 1.55 | 4.2 | 1.52 |
| What metal can you melt in the palm of your hand? | Gallium | STEM | Categorical | 5.22 | 1.26 | 4.07 | 2.02 |
| What method of underwater detection is short for sound navigation and ranging? | Sonar | STEM | Categorical | 3.27 | 1.75 | 4.33 | 1.53 |
| What nation produces two thirds of the world's vanilla? | Madagascar | Geography | Categorical | 4.06 | 1.75 | 4.17 | 1.34 |
| What nation was bounced from the Organization of American States in 1962? | Cuba | History | Categorical | 3.28 | 1.74 | 4 | 1.76 |
| What nutty legume accounts for one sixth of the world's vegetable oil production? | The peanut | General | Categorical | 4.2 | 1.86 | 4.33 | 1.15 |
| What physicist's last words were not understood because his nurse did not speak German? | Albert Einstein | History | Categorical | 4.07 | 1.71 | 3.11 | 2.09 |
| What planet is closest in size to our moon? | Mercury | STEM | Categorical | 4.56 | 1.67 | 4.21 | 1.42 |
| What popular drink did a Dutch medical professor produce in his lab when trying to come up with a blood cleanser that could be sold in drugstores? | Gin | STEM | Categorical | 4 | 2.12 | 4.19 | 1.52 |
| What popular food was invented in the 1800s and sold as a diarrhea cure? | Ketchup | History | Categorical | 4 | 1.7 | 5.07 | 0.8 |
| What sort of creature appeared in "Free Willy"? | Whale | Pop Culture | Categorical | 3.25 | 2.25 | 2.75 | 1.75 |
| What spiny venous fish, common in home aquariums, has become an invasive species in the Caribbean Sea and U.S. Atlantic coastal waters? | Lionfish | Animals | Categorical | 4.06 | 1.73 | 4.09 | 1.22 |
| What television host quipped at his 1990 wedding, 'The answer is ... yes'? | Alex Trebek | Pop Culture | Categorical | 2.6 | 1.82 | 3 | 1.91 |
| What term is short for 'binary digit'? | Bit | STEM | Categorical | 3.56 | 1.46 | 3.45 | 1.57 |
| What type of animal produces a material called gossamer? | Spider | Animals | Categorical | 3.29 | 1.8 | 4.55 | 1.57 |
| What type of elephant has the biggest ears? | African | Animals | Categorical | 3.29 | 1.23 | 3.68 | 1.95 |
| What type of food is a 'hen of the woods'? | A mushroom | General | Categorical | 3.63 | 1.79 | 3.71 | 1.68 |
| What unit of electrical power is equal to one joule per second? | The Watt | STEM | Categorical | 3.5 | 1.86 | 3.5 | 1.21 |
| What was featured in the first TV commercial advertising a toy? | Mr. Potato Head | Pop Culture | Categorical | 3.57 | 1.6 | 3.19 | 1.64 |
| What was Robert Redford's first movie? | War Hunt | Pop Culture | Categorical | 2.36 | 1.75 | 3.73 | 1.91 |
| What was the favourite food of the Teenage Mutant Ninja Turtles? | Pizza | Pop Culture | Categorical | 2.3 | 1.7 | 2.14 | 1.52 |
| What was the first commercially manufactured breakfast cereal? | Shredded Wheat | History | Categorical | 4.54 | 1.5 | 4.12 | 1.5 |
| What was the first organ successfully transplanted from a cadaver to a live person? | A kidney | STEM | Categorical | 4.37 | 1.89 | 5 | 1.6 |
| What was the first planet to be discovered using the telescope, in 1781? | Uranus | History | Categorical | 3.61 | 1.97 | 4.29 | 1.96 |
| When referring to an establishment that sells alcoholic drinks, what is the word 'pub' short for? | Public House | General | Categorical | 4.56 | 1.21 | 4.14 | 2.12 |
| When there are two full moons in the same month, what is the second called? | Blue Moon | STEM | Categorical | 4 | 1.83 | 4.5 | 1.96 |
| When was the website Facebook launched? | 2004 | Pop Culture | Numeric | 3.35 | 1.79 | 3.21 | 1.58 |
| Where did the pineapple plant originate? | South America | Geography | Categorical | 4 | 1.76 | 4.71 | 1.33 |
| Where is the world's largest supply of fresh water? | Brazil | Geography | Categorical | 4.38 | 1.82 | 4.54 | 1.33 |
| Where would you find the Sea of Tranquility? | The moon | STEM | Categorical | 4.11 | 1.82 | 3.67 | 1.63 |
| Which 90's movie soundtrack is the best-selling soundtrack of all time? | The Bodyguard | Pop Culture | Categorical | 4.28 | 1.77 | 3.8 | 1.88 |
| Which American author wrote the non-fiction novel 'In Cold Blood'? | Truman Capote | Pop Culture | Categorical | 2.96 | 1.74 | 4 | 1.41 |
| Which anti-impotence treatment became the 90s fastest-selling prescription drug? | Viagra | History | Categorical | 4.47 | 1.33 | 4.33 | 2.08 |
| Which atmospheric gas is the most common in the Earth's atmosphere? | Nitrogen | STEM | Categorical | 3.79 | 1.42 | 4.92 | 1.44 |
| Which Buzz appeared in Toy Story? | Lightyear | Pop Culture | Categorical | 2.2 | 1.3 | 2.61 | 1.54 |
| Which Canadian Prime Minister was elected in 2015? | Justin Trudeau | General | Categorical | 2.75 | 1.71 | 2.5 | 0.71 |
| Which city had the most restaurants per capita in Canada? | Montreal | Geography | Categorical | 5.5 | 0.71 |  |  |
| Which city has the most restaurants per capita in Canada? | Montreal | Geography | Categorical | 3.73 | 1.62 | 4.19 | 1.83 |
| Which country has more tractors per capita, Canada, Iceland or Japan? | Iceland | Geography | Categorical | 4.75 | 1.42 | 4 | 1.5 |
| Which country has not fought in a war since 1814? | Sweden | General | Categorical | 4.22 | 1.52 | 4.85 | 1.57 |
| Which country is known as the Pearl of Africa? | Uganda | Geography | Categorical | 4.18 | 1.59 | 4.35 | 1.14 |
| Which famous lady participated in the opening of Walt Disney's first European theme park, outside of Paris, in 1992? | Cher | Pop Culture | Categorical | 3.6 | 1.65 | 3.05 | 1.47 |
| Which fictional character was also known as Lord Greystoke? | Tarzan | Pop Culture | Categorical | 2.35 | 1.5 | 2.13 | 0.99 |
| Which instrument did Louis Armstrong play? | Trumpet | History | Categorical | 3.93 | 1.94 | 4.29 | 1.38 |
| Which is the only mammal that can't jump? | Elephant | Animals | Categorical | 4.44 | 1.59 | 4.86 | 1.41 |
| Which ocean is off the Californian coast? | Pacific | Geography | Categorical | 3.13 | 1.96 | 5 |  |
| Which province/territory is home to Canada's tallest mountain? | Yukon | Geography | Categorical | 4.42 | 1.71 | 3.5 | 1.73 |
| Which scientist is considered the father of modern genetics? | Gregor Mendel | STEM | Categorical | 3.6 | 2.17 | 4.63 | 0.92 |
| Which shoe company did Michael Jordan famously promote? | Nike | Sports | Categorical | 4.31 | 2.22 | 4.17 | 1.94 |
| Which state is called the Empire State? | New York | General | Categorical | 4.5 | 1.29 | 4 | 2.83 |
| Which state is called the Golden state? | California | General | Categorical | 4.21 | 1.72 |  |  |
| Which type of semi aquatic animal is a lutra-lutra? | An otter | Animals | Categorical | 4.05 | 1.62 | 3.76 | 2.1 |
| Which U.S. president issued the Emancipation Proclamation? | Abraham Lincoln | History | Categorical | 2.35 | 1.84 | 5.2 | 0.84 |
| Which U.S. President made the first telephone call to the moon? | Richard Nixon | History | Categorical | 3.74 | 1.76 | 4.29 | 1.38 |
| Which US President was inaugurated in 1933? | Franklin Roosevelt | History | Categorical | 3.15 | 1.73 | 2.88 | 1.81 |
| Which vitamin is also known as pantothenic acid? | B5 | STEM | Numeric | 3.48 | 1.58 | 3.56 | 1.46 |
| Which was the first nation to give women the right to vote? | New Zealand | History | Categorical | 4.55 | 1.79 | 5.79 | 0.43 |
| Who created Bugs Bunny? | Tex Avery | Pop Culture | Categorical | 3.18 | 1.56 | 3.93 | 1.67 |
| Who had a 1960s No. 1 hit with 'The Lion Sleeps Tonight'? | The Tokens | Pop Culture | Categorical | 3.32 | 1.95 | 2.9 | 1.66 |
| Who had a big 80s No 1 with 'Every Breath You Take'? | The Police | Pop Culture | Categorical | 3.47 | 2 | 2.42 | 1.31 |
| Who holds the record for the most home runs in a single major league baseball season? | Barry Bonds | Sports | Categorical | 2.19 | 1.72 | 2.94 | 1.98 |
| Who invented the rabies vaccination? | Louis Pasteur | History | Categorical | 4.07 | 1.79 | 4.45 | 1.29 |
| Who is the lead singer for the rock band Guns N Roses? | Axl Rose | Pop Culture | Categorical | 2.32 | 1.09 | 2.05 | 1.31 |
| Who played Jack in Titanic? | Leonardo DiCaprio | Pop Culture | Categorical | 1.67 | 1.15 | 3.4 | 1.82 |
| Who played Rachel Green in Friends? | Jennifer Aniston | Pop Culture | Categorical | 3 | 1.94 | 2.56 | 0.89 |
| Who preceded Ronald Reagan as American president? | Jimmy Carter | History | Categorical | 3.44 | 1.69 | 3.17 | 2.14 |
| Who recorded the album Dark Side of the Moon? | Pink Floyd | Pop Culture | Categorical | 3.06 | 1.39 | 2.27 | 1.79 |
| Who said, 'Money is like an arm or a leg, use it or lose it'? | Henry Ford | General | Categorical | 3.88 | 1.5 | 3.2 | 1.61 |
| Who said: 'I'm the president of the United States and I'm not going to eat any more broccoli'? | George Bush | History | Categorical | 3.64 | 1.91 | 3.63 | 1.3 |
| Who succeeded Richard Nixon as President of the USA in 1974? | Ford | History | Categorical | 2.83 | 1.85 | 3.33 | 0.82 |
| Who was prime minister before Stephen Harper? | Paul Martin | General | Categorical | 2.94 | 1.55 | 4 | 1.76 |
| Who was the first boxer to regain a lost world heavyweight title? | Floyd Patterson | Sports | Categorical | 2.91 | 1.72 | 3.24 | 1.76 |
| Who was the first democratically-elected president of Russia? | Boris Yeltsin | History | Categorical | 2.65 | 1.37 | 3.56 | 1.9 |
| Who was the first female Prime Minister of a European country? | Margaret Thatcher | History | Categorical | 3.41 | 2 | 5.13 | 0.99 |
| Who was the first woman to be inducted into the Rock and Roll Hall of Fame? | Aretha Franklin | History | Categorical | 3.25 | 1.98 | 4.6 | 1.64 |
| Who was the Greek god of wine? | Dionysus | General | Categorical | 4.35 | 1.93 | 3.63 | 1.85 |
| Who was the star of the popular 80s crime drama Magnum P.I.? | Tom Selleck | Pop Culture | Categorical | 2.62 | 1.75 | 2.18 | 1.47 |
| Who won more Academy Awards in his lifetime than any other person? | Walt Disney | General | Categorical | 3.61 | 2.1 | 4.13 | 1.94 |
| Who wrote the novel East Of Eden? | John Steinbeck | Pop Culture | Categorical | 3.39 | 2.08 | 4.18 | 1.6 |
| Who wrote the novel The Great Gatsby? | F Scott Fitzgerald. | Pop Culture | Categorical | 3.5 | 1.29 | 4.83 | 1.17 |
| With twelve Oscar nominations and three wins, who is the most nominated male actor in Academy Awards history? | Jack Nicholson | Pop Culture | Categorical | 3.73 | 2.14 | 3.82 | 1.78 |
| With which sport is Babe Ruth associated? | Baseball | Sports | Categorical | 3.4 | 2.41 | 2.05 | 1.31 |

